# Hepatocyte PPARγ contributes to the progression of non-alcoholic steatohepatitis in male and female obese mice

**DOI:** 10.1101/2022.06.06.494901

**Authors:** Samuel M. Lee, Jose Muratalla, Saman Karimi, Alberto Diaz-Ruiz, Maria Dolores Frutos, Grace Guzman, Bruno Ramos-Molina, Jose Cordoba-Chacon

## Abstract

**Background & Aims:** Non-alcoholic steatohepatitis (NASH) is associated with obesity and increased expression of hepatic peroxisome proliferator-activated receptor γ (PPARγ) in humans. Although we previously showed that the expression of PPARγ in hepatocytes contributes to the development NASH in lean mice, the relevance of hepatocyte PPARγ in the development of NASH associated with obesity is still poorly understood.

**Methods:** Hepatocyte PPARγ was knocked out (Pparg^ΔHep^) after the development of high-fat diet-induced obesity in male and female mice and before NASH was induced with a high fat, cholesterol and fructose (HFCF) diet. We assessed the effect of the diets and Pparg^ΔHep^ on body composition and glucose homeostasis, as well as on the liver pathology, gene expression, and metabolome. In addition, liver biopsies from a cohort of 102 bariatric surgery patients were assessed for liver histology and gene expression.

**Results:** PPARγ expression, specifically PPARγ2, is mostly derived from hepatocytes and increased by high fat diets. Pparg^ΔHep^ reduced HFCF-induced NASH progression without altering steatosis. Interestingly, Pparg^ΔHep^ reduced the expression of key genes involved in hepatic fibrosis in HFCF-fed male and female mice, and collagen- stained fibrotic area in the liver of HFCF-fed male mice. In addition, transcriptomic and metabolomic data suggested that HFCF-diet regulated hepatic amino acid metabolism in a hepatocyte PPARγ-dependent manner. Specifically, Pparg^ΔHep^ increased betaine-homocysteine methyltransferase expression and reduced homocysteine levels in HFCF- fed male mice. In a cohort of 102 bariatric surgery patients, 16 cases of NASH were associated with increased insulin resistance and hepatic PPARγ expression.

**Conclusions:** Hepatocyte PPARγ expression associated with obesity could regulate methionine metabolism and the progression of fibrosis in NASH.

## Introduction

Non-alcoholic fatty liver disease (NAFLD) is emerging as the primary cause of chronic liver disease and has a worldwide prevalence above 25% [1, 2]. NAFLD is characterized by liver steatosis: accumulation of fat in >5% of hepatic parenchyma. Although the early stages of NAFLD are associated with low to moderate levels of steatosis NAFLD can progress into non–alcoholic steatohepatitis (NASH) which is characterized by severe steatosis, hepatocyte ballooning, and lobular inflammation with or without fibrosis [3]. Obesity and type 2 diabetes (T2DM) are major risk factors that accelerate the development of steatosis and the progression of NAFLD in humans [4]. The development of insulin resistance in obese patients increases white adipose tissue lipolysis causing a redistribution of excess non-esterified fatty acids (NEFA) from adipose tissue to the liver. Furthermore, increased postprandial carbohydrates and insulin levels enhance hepatic de novo lipogenesis (DNL). The re-esterification of fatty acids derived from adipose tissue lipolysis, and those synthetized by DNL, increases the level of triglycerides (TG) and other lipids in hepatocytes and promotes steatosis and hepatocyte ballooning [4]. An excessive accumulation of lipids in hepatocytes is associated with mitochondrial and endoplasmic reticulum dysfunction, an increased level of reactive oxygen species, and the induction of lipotoxicity. Consequently, hepatocytes release damage-associated molecular patterns that activate non-parenchymal cells and promote inflammation and fibrosis [3, 5]. To date, NAFLD/NASH imposes a significant burden on the healthcare system, which is exacerbated by the fact that there is still no FDA-approved pharmacological treatment specifically for NAFLD/NASH [1, 3]. As such, further research is required to understand the molecular mechanisms that promote the development of this disease.

Peroxisome proliferator-activated receptor gamma (PPARγ) is a well-known nuclear receptor in adipocytes that regulates the expression of genes involved in adipogenesis and the metabolism of both glucose and lipids. [6]. PPARγ is also expressed in other tissues where it regulates multiple genes associated with metabolism, inflammation, food intake, and insulin sensitivity [7, 8]. Although PPARγ is expressed at low levels in the livers of lean patients [9], its expression is significantly increased in the livers of patients with NAFLD [10–12]. Similarly, the expression of PPARγ is also increased in the livers of mouse models with NAFLD [13–23]. Our group has recently reported that the loss of hepatocyte-specific expression of PPARγ (Pparg^ΔHep^), before and after the development of diet-induced obesity in adult male mice, reduced diet-induced liver steatosis [14, 16] . Furthermore, we have shown that Pparg^ΔHep^ reduced the progression of steatohepatitis in mice fed a methionine and choline deficient diet [15], and NASH in mice fed a high fat, fructose and cholesterol (HFCF) diet [13]. While these findings are relevant, a methionine and choline deficient diet reduces body weight and insulin resistance, and 24 weeks of the HFCF diet failed to promote obesity or insulin resistance that regular high fat diets (HF, HFD) do [13, 14, 24]. Considering that obesity and T2DM are major risk factors for the development and progression of NAFLD, it is critical to determine if the expression of PPARγ in hepatocytes promotes NASH in mice with pre-established diet-induced obesity. To this end, we measured hepatic PPARγ expression in 102 bariatric surgery patients after scoring NAFLD in liver biopsies, and induced Pparg^ΔHep^ in male and female mice with pre-established diet-induced obesity and insulin resistance before feeding the HFCF diet to promote NASH.

## Results

### PPARγ and PPARγ-target genes expression was increased in the livers of obese patients with NASH

Liver biopsies were obtained from 102 obese patients undergoing bariatric surgery (25 men/77 women). Histological analysis of the liver biopsies identified that 30 patients (1 men/ 29 women, average age 44.93 years) did not have liver steatosis (No NAFLD), 56 patients (21 men/35 women, average age 47 years) had NAFLD, and 16 patients (3 men/13 women, average age 48.12 years) had NASH. None of these patients had fibrosis suggesting that this cohort of patients had early stages of NAFLD/NASH. Whereas the NAFLD stage was not dependent on body mass index for the study patients (**Figure 1A**), patients with NASH had higher glycated hemoglobin A1c (HbA1c), homeostatic model assessment for insulin resistance (HOMA-IR), and alanine transaminase (ALT) than patients with obesity without NASH (**Figure 1B-D**). In these liver samples, we measured the expression of *PPARγ*, and the PPARγ-targets: fatty acid translocase (*CD36*), and Cell Death Inducing DFFA Like Effector A (*CIDEA*). Interestingly, patients with NAFLD and NASH showed an increased hepatic expression of *PPARγ* and *CD36,* but not *CIDEA* (**Figure 1E**).

**Figure 1:**
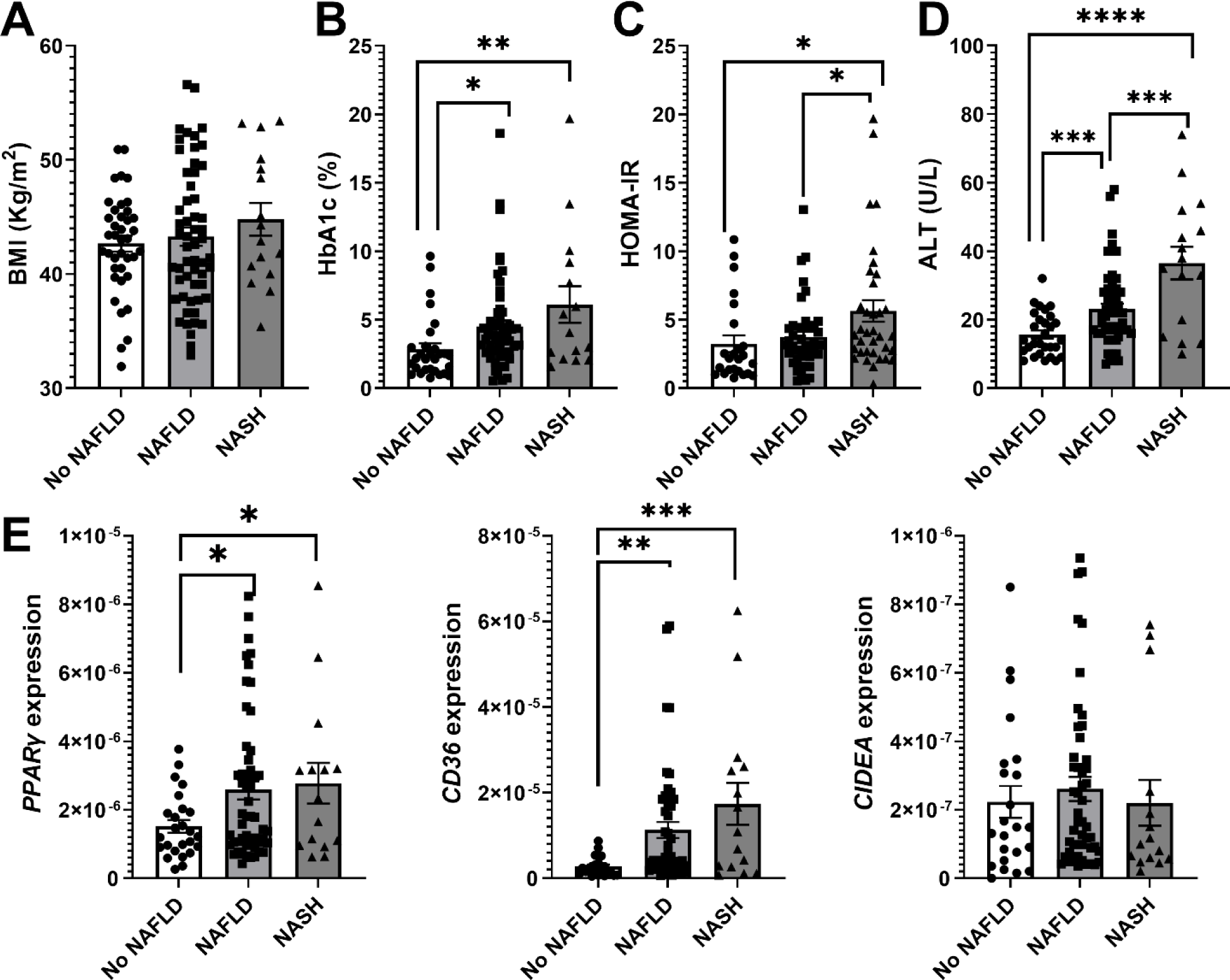
Hepatocyte PPARγ expression is increased in obese patients with NAFLD and insulin resistance. A) Body mass index (BMI, kg/m^2^), B) HbA1c, C) HOMA-IR, D) serum ALT levels, and E) expression of hepatic *PPARγ*, *CD36*, and *CIDEA*. 30 patients did not have NAFLD, 55 patients had NAFLD, and 16 patients had NASH. Asterisks indicate significant differences between groups. *, p<0.05; **, p<0.01; ***, p<0,001; ****, p<0,0001.

### PPARγ and PPARγ-target genes were increased in the livers of diet-induced obese mice that were fed a high-fat, cholesterol, and fructose diet

The expression of PPARγ in the liver was increased in male and female mice fed an HF and HFCF diet (**Figure 2A**). Consistent with our previous reports [13–16, 34], Pparg^ΔHep^ mice showed a dramatic reduction in the mRNA levels of hepatic *PPARγ* (**Figure 2A**), indicating that a significant proportion of hepatic PPARγ expression in the liver is derived from hepatocytes. High-fat diets mediated the upregulation only of hepatic PPARγ2. Independent of diet, Pparg^ΔHep^ reduced the expression of PPARγ2 in male and female mice, whereas PPARγ1 was reduced only in female mice (**Figure 2B,C**). Overall, these results suggest that PPARγ1 is primarily expressed in non- parenchymal cells in male mice and in non-parenchymal cells and hepatocytes in female mice, whereas PPARγ2 is mostly expressed in hepatocytes of male and female mice and regulated by diet. Furthermore, the hepatic expression of PPARγ targets: cell death activator A and C (*Cidea* & *Cidec*), monoacylglycerol O-acyltransferase 1 (*Mogat1*) and perilipin 4 (*Plin4*) [35, 36] were increased by HFD in male and female mice in a PPARγ-dependent manner (**Figure 2D**) suggesting that they could be under the control of PPARγ2 in hepatocytes.

**Figure 2:**
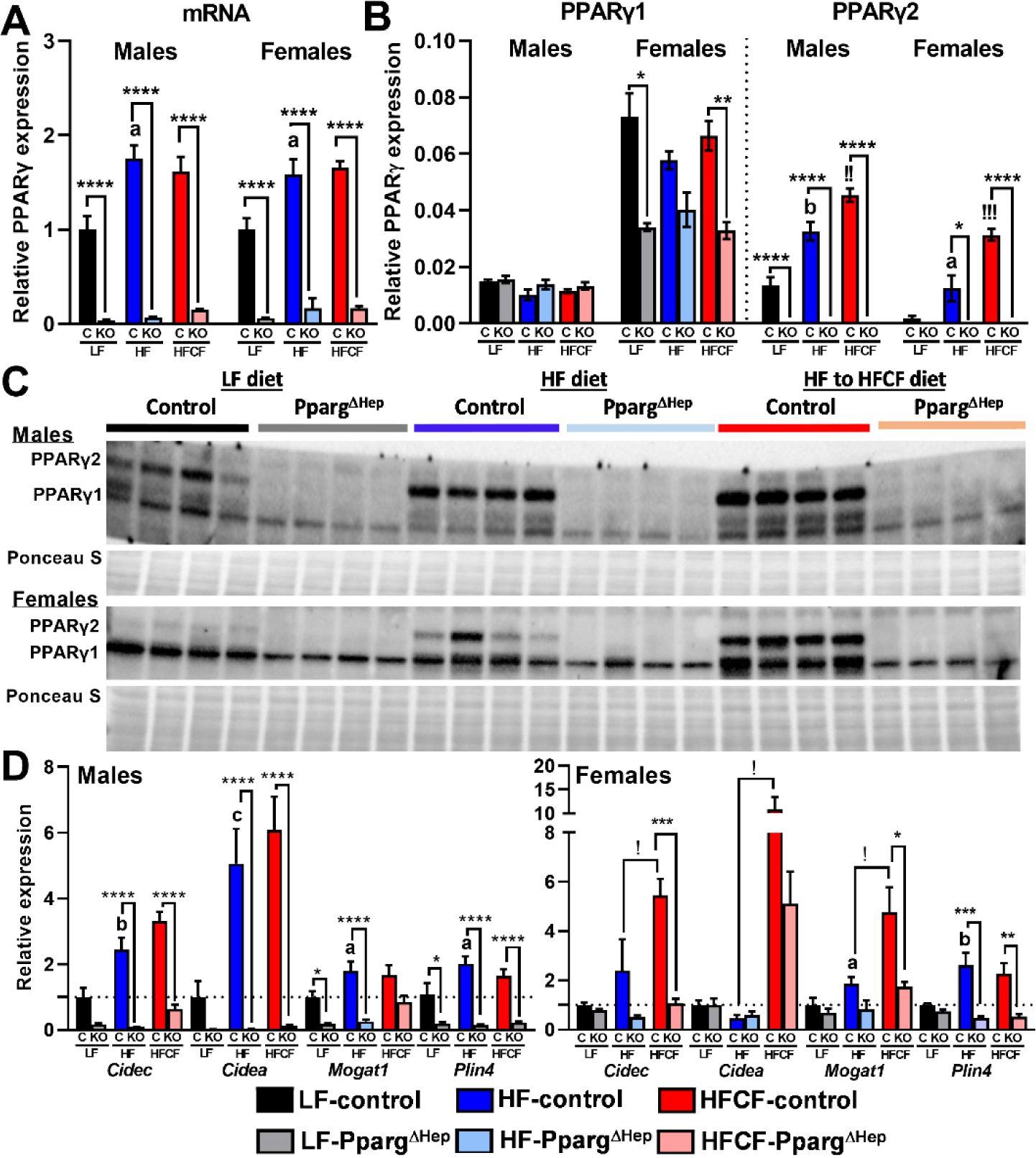
Hepatocyte PPARγ expression is increased in male and female mice with diet-induced obesity and NASH. A) PPARγ expression, B) PPARγ1 and PPARγ2 expression, C) blots of PPARγ1 and PPARγ2 bands, and Ponceau staining, and D) *Cidec, Cidea, Mogat1*, and *Plin4* expression in the liver of male and female control and Pparg^ΔHep^ mice fed a low fat diet (LF, LFD), high fat diet (HF, HFD), or high fat, cholesterol and fructose diet (HFCF). Lower letters indicate significant difference between LFD- and HFD-fed control mice. Exclamation marks indicate significant difference between HFD- and HFCF-fed control mice. Asterisks indicate significant differences between control and Pparg^ΔHep^ mice within diet. a, !, *, p < 0.05; b, !!, **, !!, p < 0.01; c, !!!, ***, p<0,001; d, ****, p< 0.0001 n = 4–9 mice/group.

### High-fat, cholesterol, and fructose diet had differential effects on adiposity and glucose homeostasis, independent of hepatocyte PPARγ expression

Next, we assessed if HF, HFCF diet or Pparg^ΔHep^ had an impact on adiposity, insulin resistance, and plasma lipids in mice with pre-established HFD-induced obesity. As expected, HFD increased body weight and adiposity in both control and Pparg^ΔHep^ mice compared to their low fat (LF/LFD)-fed male and female counterparts (**Figure 3A,B**). This was mostly due to an increase in whole-body fat mass rather than lean mass. Interestingly, when diet- induced obese mice were switched to the HFCF diet, there was a significant reduction in body weight, and fat mass in control and Pparg^ΔHep^ male mice, albeit this weight reduction was not so evident in female mice (**Figure 3A,B**). The negative effect of the HFCF diet on adiposity was confirmed by the reduced amount of individual white adipose tissue weight in both male and female mice (**Figure 3C,D**). In addition, HFD increased the area under the curve (AUC) of the glucose tolerance test (GTT) in male and female mice, and the insulin tolerance test (ITT) in male mice (**Figure 4**). Conversely, HFCF diet trended to improve the AUC of GTT in male mice, AUC of ITT in Pparg^ΔHep^ male mice, AUC of GTT in control and Pparg^ΔHep^ female mice, and reduced insulin levels in male mice (**Figure 4B-D**). Finally, we found that HFD decreased NEFA levels in control male mice and increased cholesterol levels in control male and female mice. Of note, although HFCF diet has a high concentration of cholesterol (2% w/w), this diet increased plasma cholesterol only in male mice (**Table 1**). Interestingly, Pparg^ΔHep^ increased SC and RP fat, plasma TG and decreased cholesterol in HFD-fed control male mice, without any significant effect in HFCF- fed mice.

**Figure 3:**
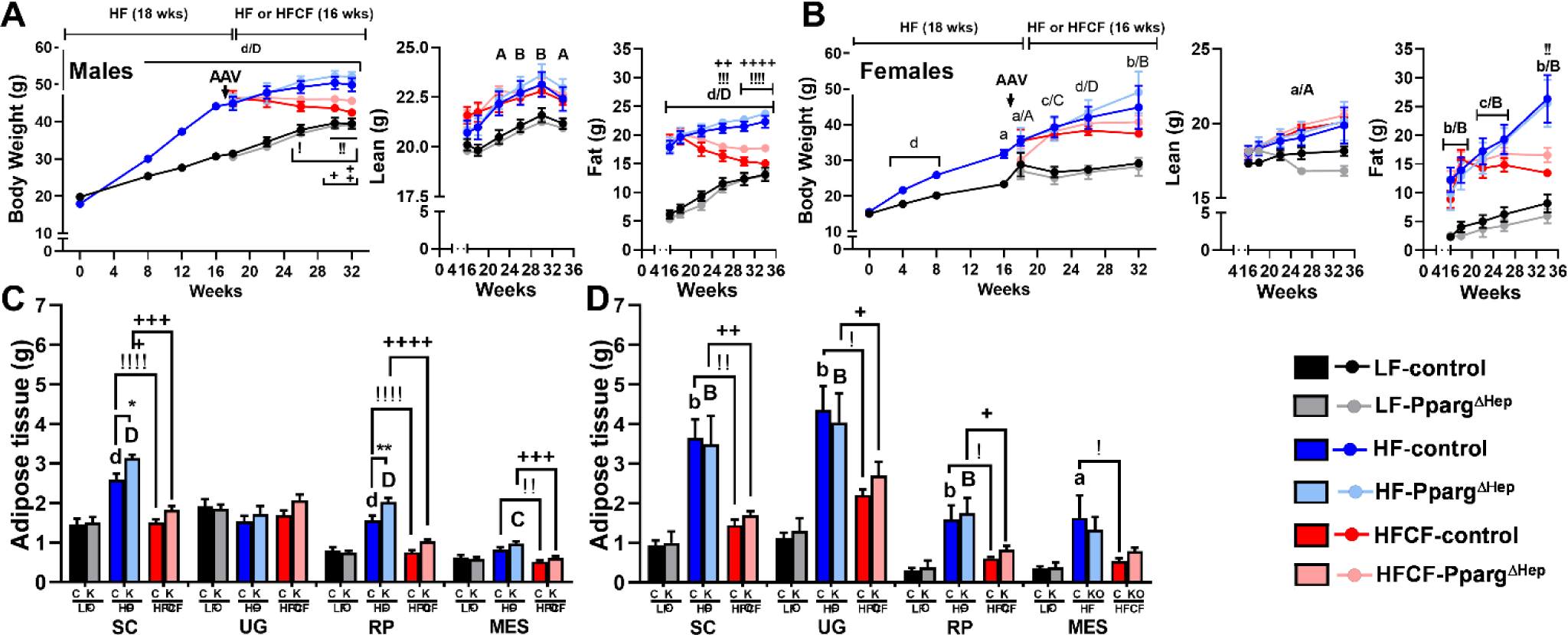
**HFCF diet reduces adiposity in diet-induced obese mice**. Body weight and whole body lean and fat mass in male (A) and female (B) mice. Weight of white adipose subdepot in male (C) and female (D) mice. Subcutanous (SC), urogenitial (UG), retroperitoneal (RP) and mesenteric (MES) adipose tissue. Lower letters indicate significant difference between LF- and HF-fed control mice. Capital letters indicate significant difference between LF- and HF-fed Pparg^ΔHep^ mice. Exclamation marks indicate significant difference between HF- and HFCF-fed control mice. Plus signs indicate significant difference between HF- and HFCF-fed Pparg^ΔHep^ mice. Asterisks indicate significant differences between control and Pparg^ΔHep^ mice within diet. a,A,!,+, *, p < 0.05; b,B,!!, ++,**, p < 0.01; c,C, p<0,001; d,D,!!!!,++++, ****, p< 0.0001 n = 4–9 mice/group.

**Figure 4:**
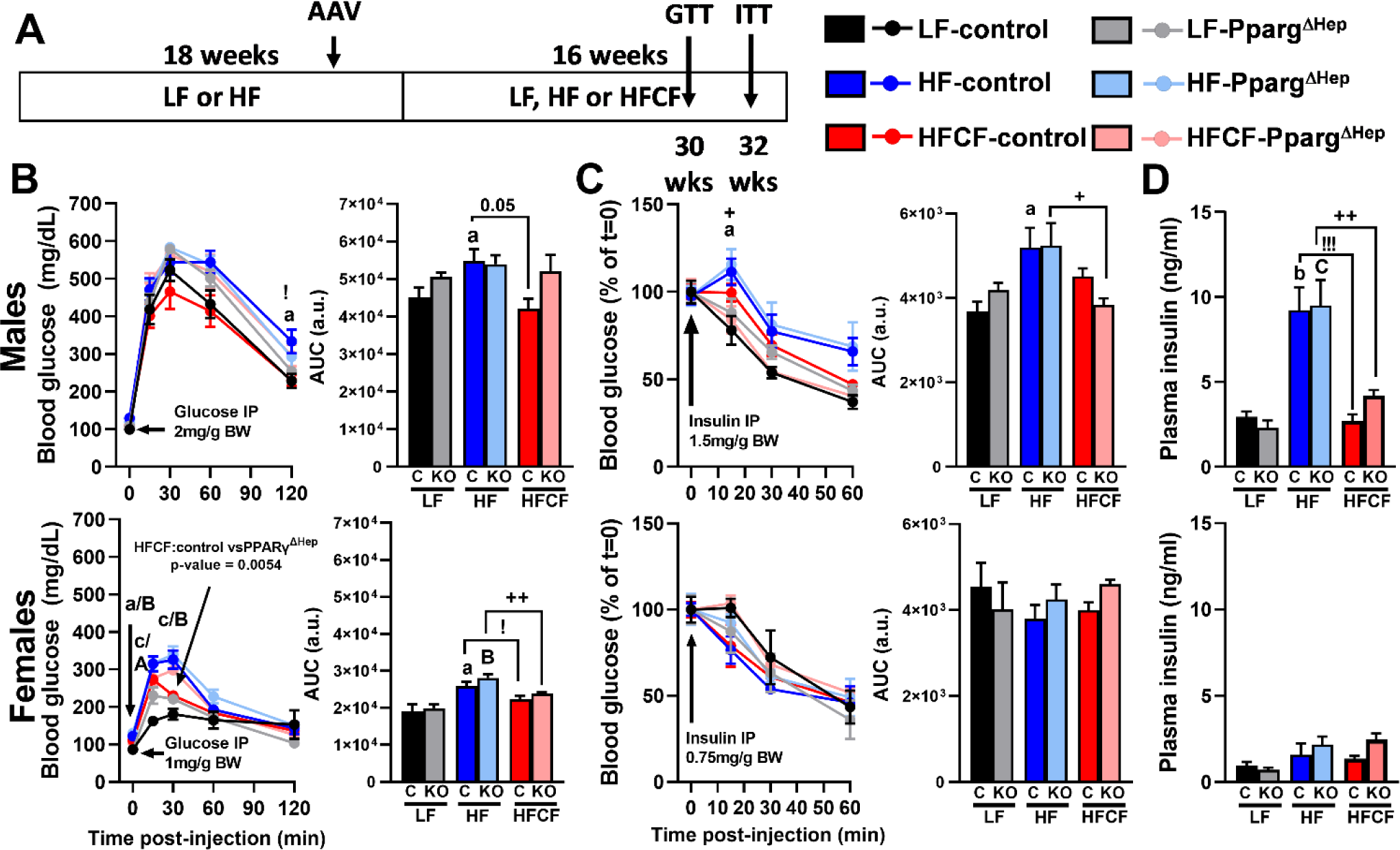
**HFCF diet improves glucose homeostasis in diet-induced obese mice**. A) Diagram with representation of GTT and ITT. B) GTT, C) ITT, and D) plasma insulin levels in male mice. E) GTT, F) ITT, and G) plasma insulin levels in female mice. GTT, glucose: 2 mg/g body weight ip in male mice, and 1 mg/g body weight ip in female mice, after 30 weeks of diet, in mice fasted overnight. ITT, insulin: 1.5 mU/g body weight ip in male mice, and 0.75 mU/g body weight ip in female mice, after 32 weeks of diet, in mice after a 4h food withdrawal at 0700h. Lower letters indicate significant difference between LF- and HF-fed control mice. Capital letters indicate significant difference between LF- and HF-fed Pparg^ΔHep^ mice. Exclamation marks indicate significant difference between HF- and HFCF-fed control mice. Plus signs indicate significant difference between HF- and HFCF-fed Pparg^ΔHep^ mice. a,A,!,+, p < 0.05; b,B, ++, p < 0.01; c,C,!!! p<0,001; n = 4–9 mice/group. AUC, area under the curve. BW, body weight. IP, intraperitoneal.

**Table 1.**
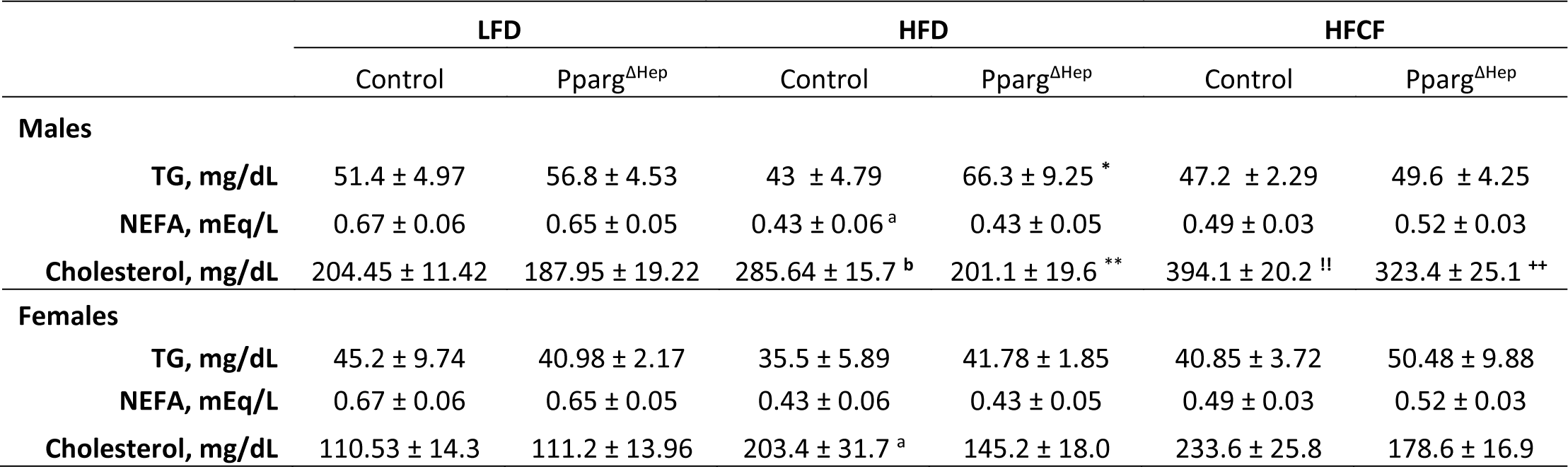
Effect of diet and Pparg^ΔHep^ in the levels of plasma lipids. Plasma TG, NEFA, and cholesterol levels were measured at the end of the study in both male and female mice fed a LFD, HFD, or HFCF diet. Lower letters indicate significant difference between LFD- and HFD-fed control mice. Exclamation marks indicate significant difference between HFD- and HFCF-fed control mice. Plus signs indicate significant differences between HFD- and HFCF-fed Pparg^ΔHep^ mice. Asterisks indicate significant difference between control and Pparg^ΔHep^ mice within diet. a, *, p < .05; b, !!, ++, p < .01 n = 4–9 mice/group.

|  | LFD |  | HFD |  | HFCF |  |
| --- | --- | --- | --- | --- | --- | --- |
|  | Control | Pparg<sup>ΔHep</sup> | Control | Pparg<sup>ΔHep</sup> | Control | Pparg<sup>ΔHep</sup> |
| **Males** |  |  |  |  |  |  |
| TG, mg/dL | 51.4 ± 4.97 | 56.8 ± 4.53 | 43 ± 4.79 | 66.3 ± 9.25 * | 47.2 ± 2.29 | 49.6 ± 4.25 |
| NEFA, mEq/L | 0.67 ± 0.06 | 0.65 ± 0.05 | 0.43 ± 0.06 <sup>a</sup> | 0.43 ± 0.05 | 0.49 ± 0.03 | 0.52 ± 0.03 |
| Cholesterol, mg/dL | 204.45 ± 11.42 | 187.95 ± 19.22 | 285.64 ± 15.7 <sup>b</sup> | 201.1 ± 19.6 ** | 394.1 ± 20.2 <sup>!!</sup> | 323.4 ± 25.1 <sup>++</sup> |
| **Females** |  |  |  |  |  |  |
| TG, mg/dL | 45.2 ± 9.74 | 40.98 ± 2.17 | 35.5 ± 5.89 | 41.78 ± 1.85 | 40.85 ± 3.72 | 50.48 ± 9.88 |
| NEFA, mEq/L | 0.67 ± 0.06 | 0.65 ± 0.05 | 0.43 ± 0.06 | 0.43 ± 0.05 | 0.49 ± 0.03 | 0.52 ± 0.03 |
| Cholesterol, mg/dL | 110.53 ± 14.3 | 111.2 ± 13.96 | 203.4 ± 31.7 <sup>a</sup> | 145.2 ± 18.0 | 233.6 ± 25.8 | 178.6 ± 16.9 |

### Hepatocyte PPARγ contributes to the development of the hepatic phenotype in diet-induced obese mice fed a high-fat, cholesterol and fructose diet

We have previously assessed the effect of hepatocyte PPARγ in the development of steatosis before and after diet-induced obesity is established [14, 16], and before HFCF-induced NASH [13]. In this study, feeding a HFD (60% Kcal from fat) for 34 weeks did not significantly increase liver weight in male or female mice as compared to their LFD-fed littermates (**Figure 5A**). However, HFD increased steatosis in male and female control mice (**Figure 5B**). To further assess the progression of NASH in diet-induced obese male and female mice, we measured plasma ALT, assessed the histology of hematoxylin and eosin (H&E)-stained liver sections following the Kleiner score system, and measured the expression of genes involved in inflammation. HFD diet alone did not increase ALT levels but increased NAFLD activity score (NAS) due to the development of steatosis in male and female mice (**Figure 5C-E, 5G**). It should be noted that HFD did not increase inflammation in H&E-stained liver sections, nor the expression of inflammation-related genes: tumor necrosis factor alpha (*Tnfα*) and monocyte chemoattractant protein 1 (*Mcp1*) (**Figure 5F-I**). While there was a modest increase of triggering receptor expressed on myeloid cells 2 (*Trem2*) in male mice (**Figure 5F**) the low inflammatory scores by two independent pathologists suggest that this expression was not associated with significant hepatic inflammation. By contrast, HFCF diet dramatically increased liver weight in diet-induced obese male and female control mice without enhancing steatosis as compared to HFD-fed mice (**Figure 5A,B**). Moreover, HFCF diet promoted NASH (NAS>5) and increased the expression of *Tnfα*, *Mcp1*, and *Trem2* (**Figure 5A-F**). In this study, two independent pathological analyses indicated that both diet-induced obese male and female mice, fed a HFCF diet for 16 weeks, developed NASH: NAS= 5.7+/-0.3 control males and 6.4+/-0.2 control females (**Figure 5D,E**). The individual scores of steatosis, inflammation, and ballooning show that although HFD significantly induced liver steatosis, HFCF diet was required to increase ballooning and inflammation to promote NASH (**Figure 5E, G-I**).

**Figure 5:**
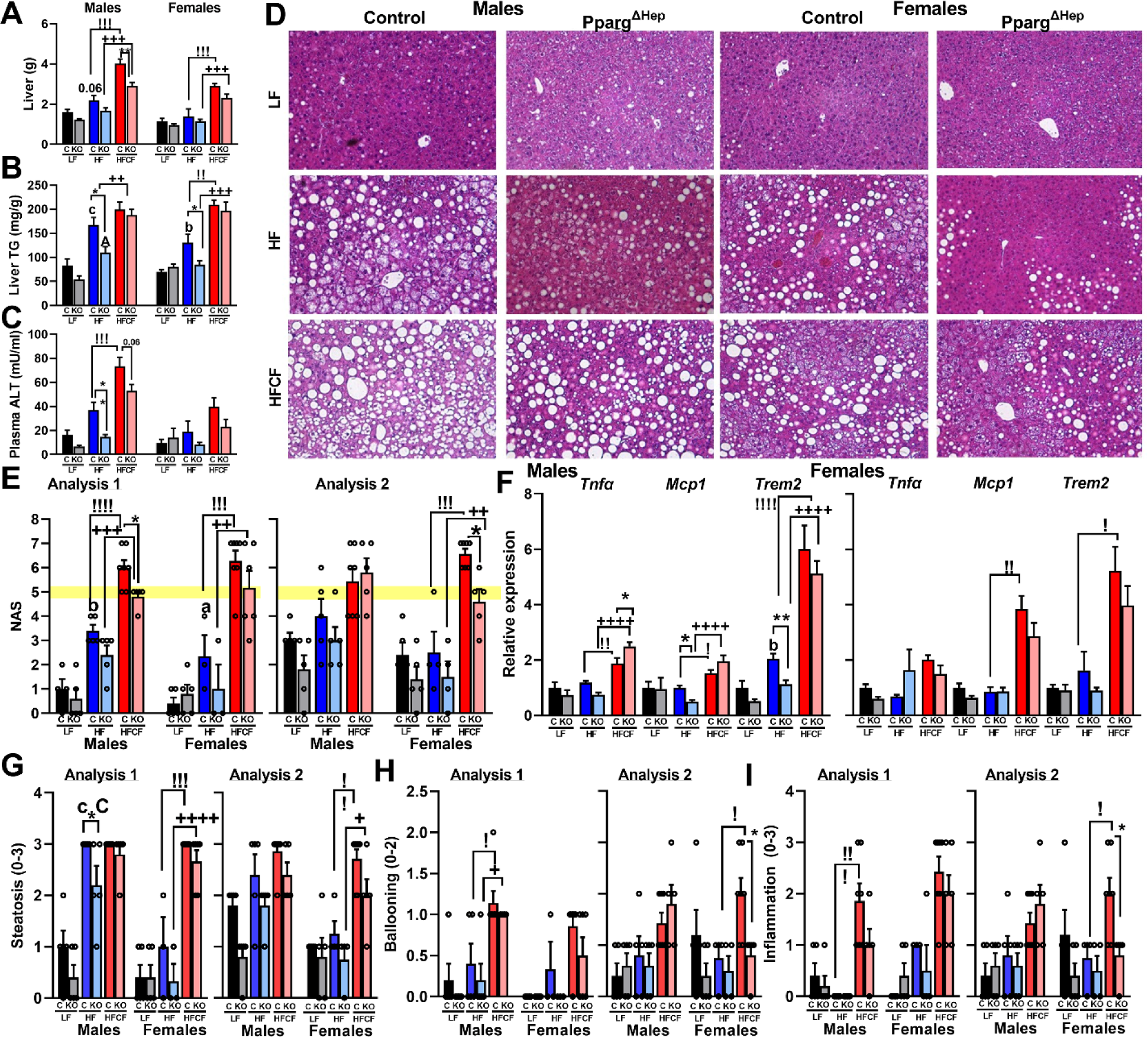
Loss of hepatocyte PPARγ expression reduces the progression of NAFLD in male and female mice. A) Liver weight, B), liver TG, C) plasma ALT, D) representative images of H&E-stained liver sections, E) NAS , F) hepatic expression of *Tnfα, Mcp1*, and *Trem2*, and individual scores of G) steatosis, H) hepatocyte ballooning, and I) inflammation of male and female control and Pparg^ΔHep^ mice fed a LF, HF, or HFCF diet. Lower letters indicate significant difference between LFD- and HFD-fed control mice. Capital letters indicate significant difference between LFD- and HFD-fed Pparg^ΔHep^ mice. Exclamation marks indicate significant difference between HFD- and HFCF-fed control mice. Plus signs indicate significant difference between HFD- and HFCF-fed Pparg^ΔHep^ mice. Asterisks indicate significant differences between control and Pparg^ΔHep^ mice within diet. a,A,!, , *, p < 0.05; b,!!, ++,**, p < 0.01; c,!!!,+++ p<0,001; d,D,!!!!,++++, p< 0.0001 n = 4–9 mice/group.

Remarkably, whereas Pparg^ΔHep^ reduced liver steatosis in HFD-fed male and female mice, and reduced plasma ALT in HFD-fed male mice, as we recently reported in HFD-induced obese mice [14, 16], Pparg^ΔHep^ reduced liver weight in male mice without reducing liver steatosis in HFCF-fed mice (**Figure 5A-C, G**). These findings suggest that the effects of hepatocyte PPARγ on liver steatosis may be restricted to mice fed an HFD diet. Although we did not observe a clear reduction in NAS in male and female Pparg^ΔHep^ mice, NAS was significantly reduced in HFCF-fed male (analysis 1) and female mice (analysis 2) by Pparg^ΔHep^ as compared to their respective HFCF-fed controls (**Figure 5E**). Of note, the proportion of mice with NASH (NAS>5) was greater in male and female control mice than in Pparg^ΔHep^ mice in both independent analyses, suggesting that Pparg^ΔHep^ may reduce the progression of NAFLD without affecting steatosis in diet-induced obese mice fed a HFCF diet.

It is known that an HFCF diet induces human-like NASH with fibrosis in mice [13, 25]. Therefore, we assessed the extent of fibrosis by assessing picrosirius red/fast green stained liver sections (**Figure 6A**) and the expression of genes associated with fibrosis. As shown in **Figure 6**, HFD did not induce fibrosis in male or female mice despite the increased expression of fibrosis-related genes: collagen 1a1 (*Col1a1*), TIMP metallopeptidase inhibitor 1 (*TIMP1*), and metallopeptidase 13 (*Mmp13*) in male mice (**Figure 6B-D**). However, HFCF diet increased the collagen-stained area detected in control male mice (**Figure 6B**), and the severity of fibrosis (bridging fibrosis, 3 and cirrhosis, 4) in both male and female control mice fed a HFCF diet (**Figure 6C**). This is supported by the increased expression of *Col1a1*, *TIMP1*, and *Mmp13* in both male and female mice fed a HFCF diet (**Figure 6D**). Of note, Pparg^ΔHep^ reduced HFCF-induced collagen-stained area detected in male mice (**Figure 6B**), the expression of *Col1a1* and *TIMP1* in both male and female HFCF-fed mice, and the expression of *Col1a1*, *TIMP1*, and *Mmp13* in HFD-fed male mice (**Figure 6D**). Altogether, these results suggest that PPARγ expression in hepatocytes may contribute to the development of fibrosis.

**Figure 6:**
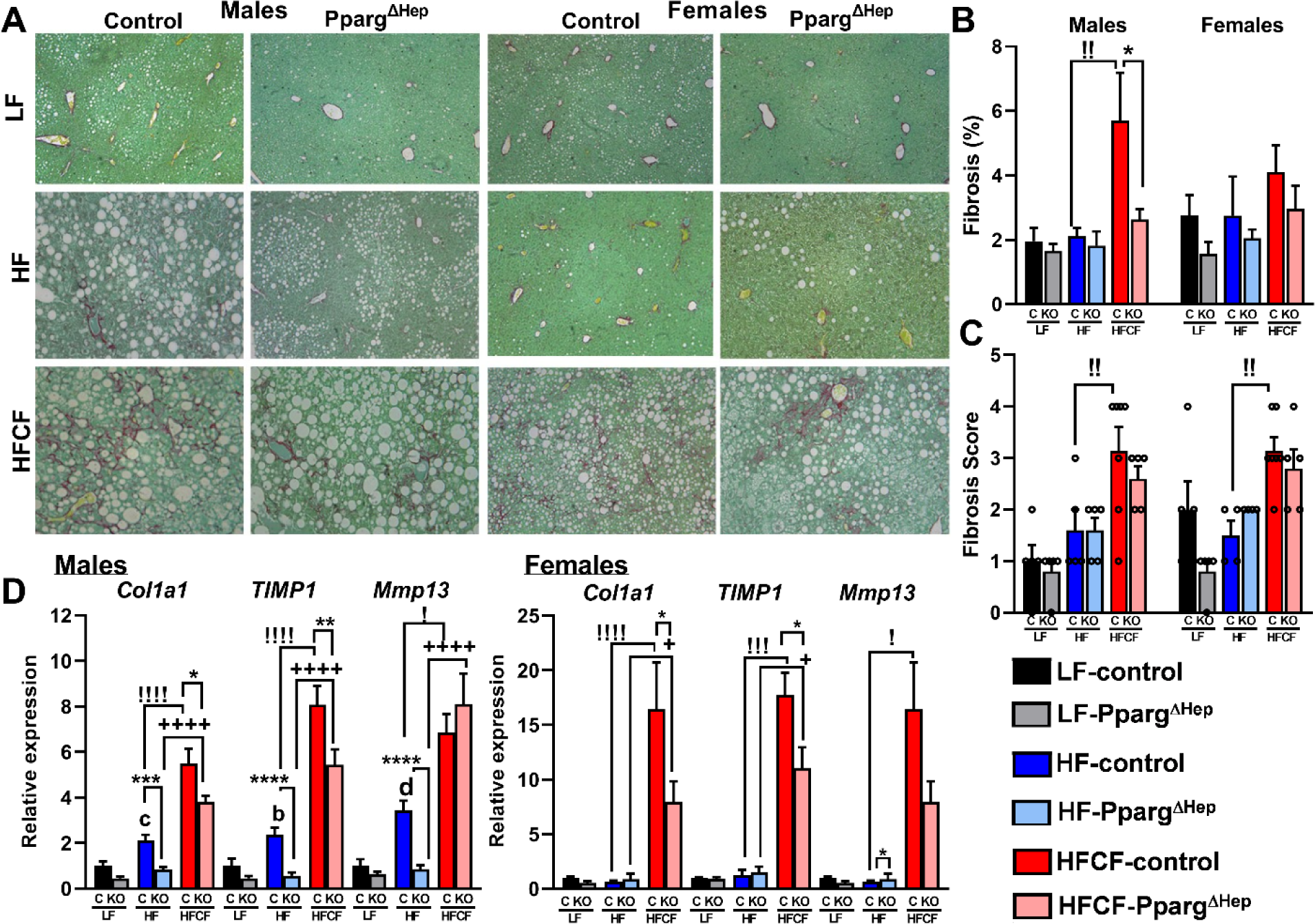
Loss of hepatocyte PPARγ expression reduces fibrosis in male mice. A) Representative images of sirius red/fast green-stained liver sections, B) percentage of collagen-stained area, C) fibrosis score plasma ALT, and D) hepatic expression of *Col1a1, TIMP1*, and *Mmp13* of male and female control and Pparg^ΔHep^ mice fed a LF, HF, or HFCF diet. Lower letters indicate significant difference between LFD- and HFD-fed control mice. Exclamation marks indicate significant difference between HFD- and HFCF-fed control mice. Plus signs indicate significant difference between HFD- and HFCF-fed Pparg^ΔHep^ mice. Asterisks indicate significant differences between control and Pparg^ΔHep^ mice within diet. !,+, *, p < 0.05; b,!!, p < 0.01; c,!!!,*** p<0,001; d,!!!!,++++,**** p< 0.0001 n = 4–9 mice/group.

### Hepatocyte PPARγ negatively regulates methionine metabolism in diet-induced obese mice fed a high-fat, cholesterol, and fructose diet

In order to shed light on the potential molecular mechanisms that hepatocyte PPARγ regulates to promote NASH, we performed a transcriptomic analysis by RNAseq on the livers of diet-induced obese control and Pparg^ΔHep^ mice fed a HFCF diet for 16 weeks. HFCF diet altered the expression of >3000 genes in diet-induced obese male and female control mice, with a similar number of differentially expressed genes (DEG, Log2 fold change >1 or <- 1) regulated by HFCF diet in both diet-induced obese control or Pparg^ΔHep^ male and female mice (**Table 2**). The enrichment analysis showed that HFCF diet regulated gene ontology (GO) terms related to fibrosis (extracellular matrix), inflammation, and mitochondrial metabolism in both diet-induced obese male and female mice (**Supplemental Tables S4-5**). These results were similar to those of our previous RNAseq performed in liver samples of male mice fed the HFCF diet for 24 weeks [13]. In our previous study, hepatocyte PPARγ was required to mediate the strong effect of HFCF diet on hepatic transcriptome (total number of DEG). However, in this study, HFCF diet regulated a similar number of DEG in both control and Pparg^ΔHep^ mice (effect of HFCF in diet-induced obese mice, **Table 2**). Pathway enrichment analysis also displayed a similar regulation in diet-induced obese control and Pparg^ΔHep^ mice fed a HFCF diet (**Supplemental Tables S4-5**). However, the number of DEG regulated by Pparg^ΔHep^ in mice fed a HFD or an HFCF diet was larger in males than in their female counterparts (**Table 2, Figure 7A,B**), which likely was indicative of a sexual dimorphism of PPARγ in the liver.

**Figure 7:**
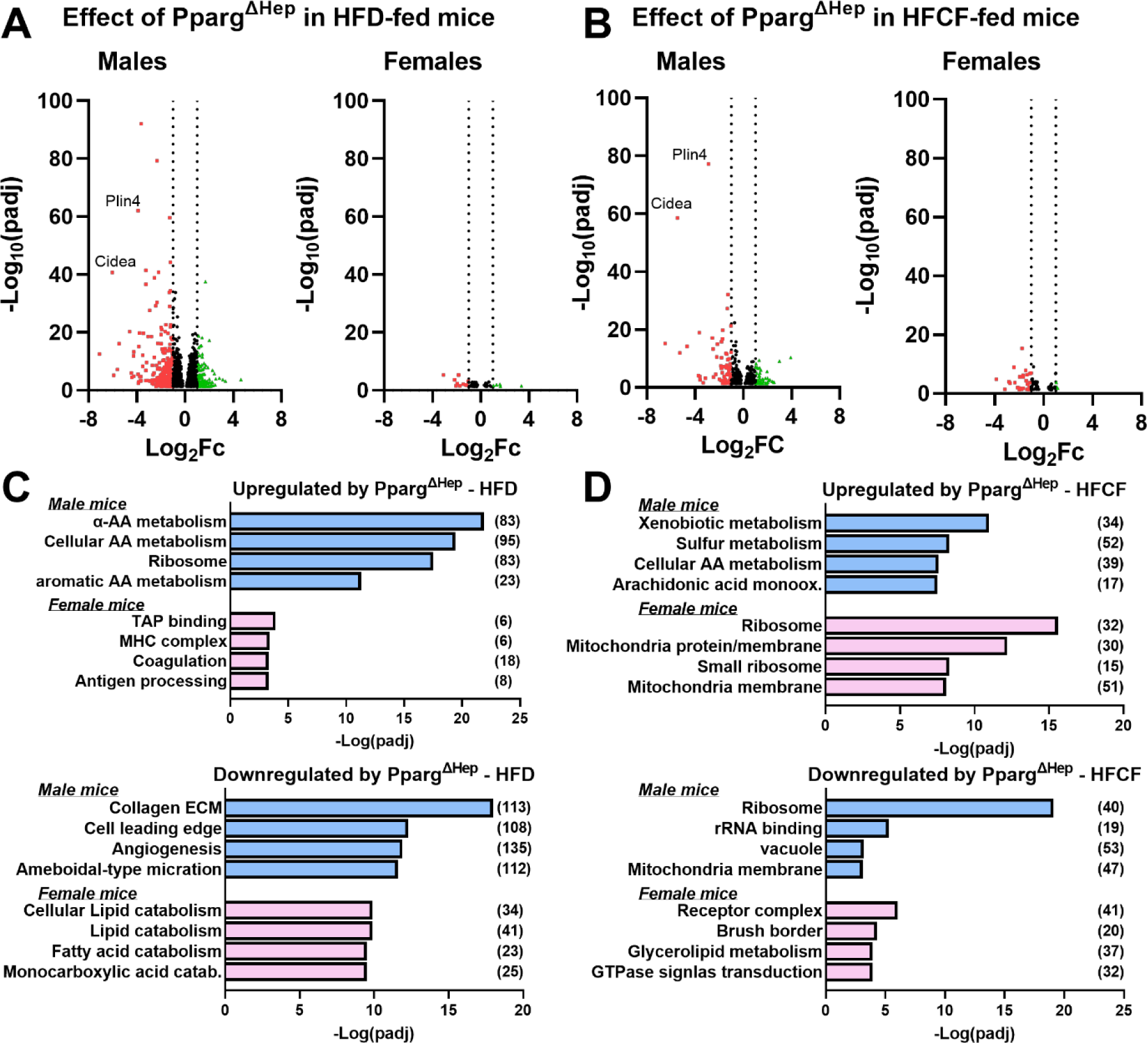
Loss of hepatocyte PPARγ expression alters the transcriptome of mice fed a HFD or a HFCF diet. Volcano plots showing the DEG (padj <0.05) by Pparg^ΔHep^ in HFD-fed (A) or HFCF-fed (B) male and female mice. Red dots show DEG with fold change (FC) < -1 (Log2). Green dots show DEG with FC>1 (Log2). Selectedgroups of genes identified by the enrichment analysis of DEG by Pparg^ΔHep^ in HFD-fed (C) or HFCF-fed (D) male and female mice. Number of DEG in these GO terms are indicated in brackets. N=4-6 mice/group.

**Table 2.** Differential expressed genes (DEG) in male and female mice. Effect of HFCF diet in diet-induced obese control and Pparg^ΔHep^ mice or effect of Pparg^ΔHep^ in HFD-fed or HFCF-fed mice. FC, fold change. n = 4–6 mice/group

|  | Effect of HFCF diet in diet-induced obese: |  |  |  | Effect of Pparg<sup>ΔHep</sup> in: |  |  |  |
| --- | --- | --- | --- | --- | --- | --- | --- | --- |
|  | Control mice |  | Pparg<sup>ΔHep</sup> mice |  | HFD-fed mice |  | HFCF-fed mice |  |
|  | Males | Females | Males | Females | Males | Females | Males | Females |
| **DEG** | 3871 | 5065 | 6287 | 3008 | 2904 | 64 | 660 | 82 |
| **Upregulated (FC [Log2]>1)** | 786 | 1773 | 1508 | 1029 | 122 | 5 | 44 | 3 |
| **Downregulated (FC[Log2] <-1)** | 121 | 283 | 210 | 238 | 309 | 11 | 68 | 27 |

Remarkably, Pparg^ΔHep^ regulated GO terms related to amino acid metabolism in both HFD- and HFCF-fed male mice (**Figure 7C,D; Supplemental Tables S6-7**). Therefore, we used liquid chromatography/mass spectrometry to measure the relative levels of a panel of 297 hydrophilic metabolites that includes metabolites related to amino acid metabolism in liver samples of control and Pparg^ΔHep^ mice. From this panel, 200-225 metabolites were identified in the livers of diet-induced obese male and female mice. HFCF diet significantly altered (p-value <0.05) the levels of 130 and 42 metabolites in the livers of diet-induced obese male and female control mice (**Figure 8A**), respectively. In this case, the effect of the HFCF diet in the liver of diet-induced obese mice seems to have a stronger effect in male: 14 metabolites with fold change < -1 (Log2FC) and 76 metabolites with fold change > 1 (Log2FC) as compared to the differentially regulated metabolites of female mice (**Figure 8A**). Furthermore, the effect of Pparg^ΔHep^ seemed to be stronger in male mice as well. Specifically, Pparg^ΔHep^ altered the levels of 55 and 51 metabolites in the livers of HFD- and HFCF-fed male mice, and only 24 and 15 metabolites in the livers of HFD- and HFCF-fed female mice, respectively (**Figure 8B,C**). These data suggest a sex-dependent effect of HFD and HFCF diet that could be PPARγ-dependent.

**Figure 8:**
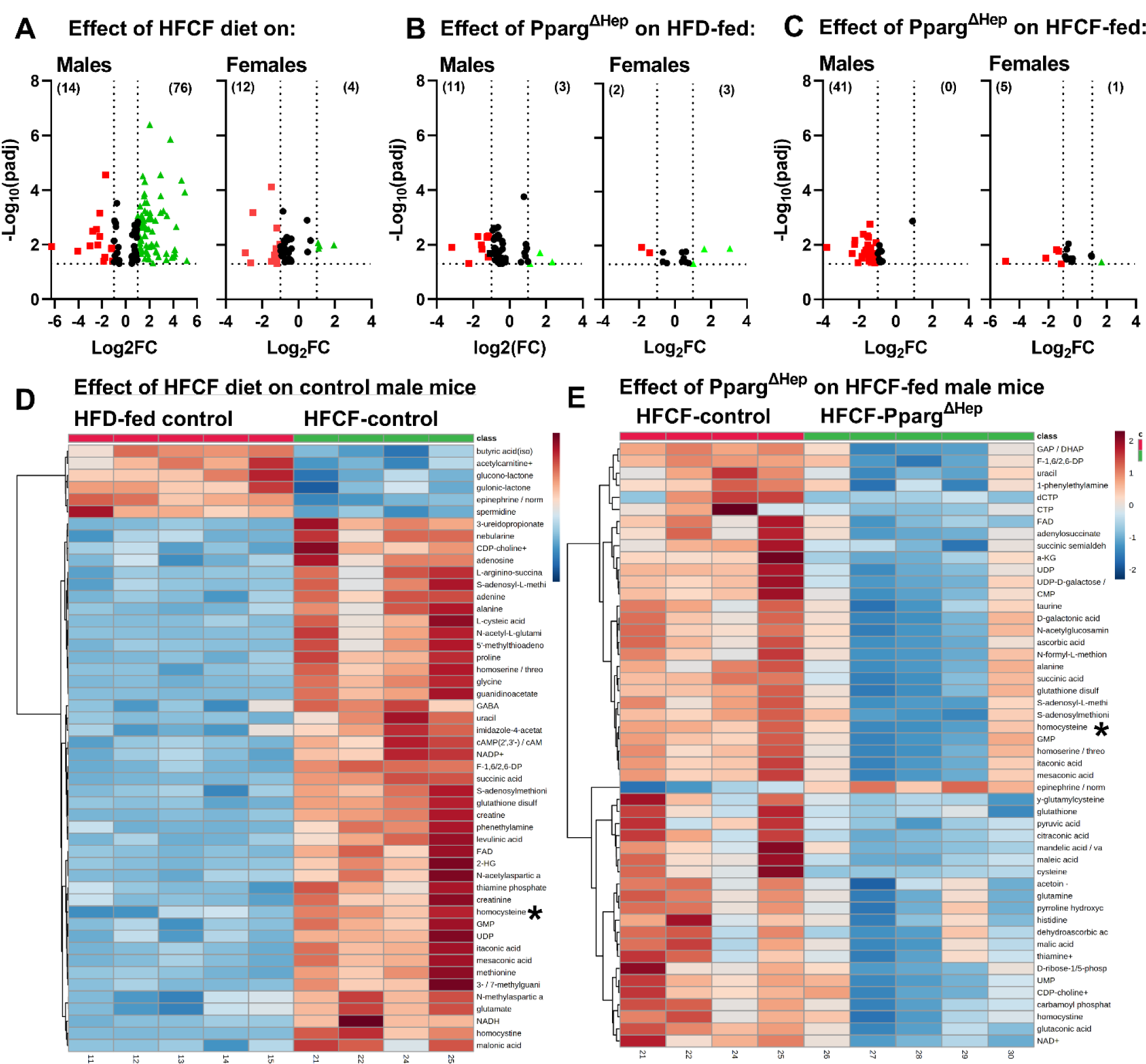
Loss of hepatocyte PPARγ expression alters the metabolome of diet-induced obese mice fed a HFD or a HFCF diet. Volcano plots showing the differentially regulated metabolites (p-value <0.05) by HFCF diet in male and female mice (A), or by Pparg^ΔHep^ in HFD-fed (B) or HFCF-fed (C) male and female mice. Red dots show differentially regulated metabolites with fold change (FC) < -1 (Log2), and green dots show DEG with FC>1 (Log2), the number of these regulated metabolites are between brackets. Heatmaps of differentially regulated metabolites between HFD- and HFCF-fed control male mice (D), or between HFCF-control and HFCF-fed Pparg^ΔHep^ mice (E). n=4-5 mice/group.

Interestingly, the heatmaps revealed the strong regulation of the metabolites mediated by HFCF diet in the liver of control male mice (**Figure 8D**) and by PPARγ in HFCF-fed male mice (**Figure 8E**). In fact, a detailed analysis of the fold changes revealed that homocysteine (Hcy) was among the metabolites differentially regulated by HFCF diet in Pparg^ΔHep^ HFCF-fed male mice. These data were similar to those published in our previous report [13] and suggested that hepatocyte PPARγ was negatively associated with the regulation of methionine metabolism in diet- induced obese mice fed a HFCF diet. Specifically, the levels of metabolites of the methionine cycle: methionine, S-adenosylmethionine (SAM), and Hcy were increased by HFCF diet in diet-induced obese male but not in female mice (**Figure 9A**). Interestingly, Pparg^ΔHep^ reduced the levels of methionine and SAM in HFCF-fed mice, and Hcy in both HFD- and HFCF-fed male mice (**Figure 9A**). Furthermore, the expression of betaine homocysteine methyltransferase (*Bhmt*), the enzyme that converts Hcy back to methionine, was increased at mRNA and protein male level in Pparg^ΔHep^ mice fed an HFCF diet (**Figure 9B-D**). Considering that excess levels of homocysteine is associated with the development of fibrosis, these results suggest that the regulation of Hcy metabolism by hepatocyte PPARγ is correlated with the progression of hepatic fibrosis.

**Figure 9:**
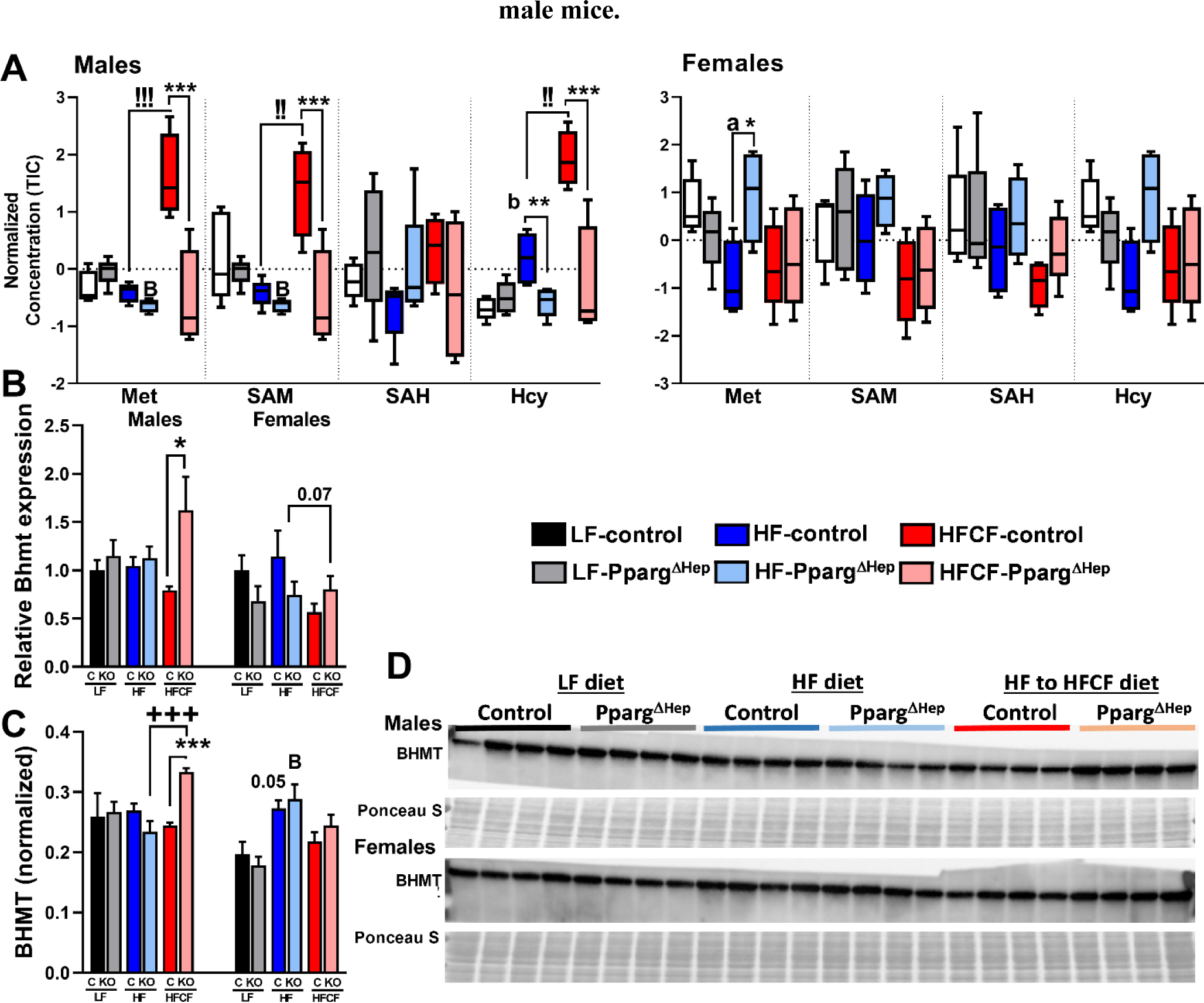
Hepatocyte PPARγ expression is needed to increase homocysteine levels in HFCF-fed male mice. A) Hepatic levels of metabolites of methionine cycle, B) mRNA expression of BHMT, and C, D) protein expression of BHMT of male and female control and Pparg^ΔHep^ mice fed a LFD, HFD, or HFCF diet. Lower letters indicate significant difference between LFD- and HFD-fed control mice. Capital letters indicate significant difference between LFD- and HFD-fed Pparg^ΔHep^ mice. Exclamation marks indicate significant difference between HFD- and HFCF-fed control mice. Plus signs indicate significant difference between HFD- and HFCF-fed Pparg^ΔHep^ mice. Asterisks indicate significant differences between control and Pparg^ΔHep^ mice within diet. a *, p < 0.05; b,B,!!,** p < 0.01; c,!!!,+++,*** p<0,001. n = 4–9 mice/group.

## Discussion

The role of hepatic PPARγ in the development of NAFLD is a subject of debate. Some review articles suggest, without strong evidence, that hepatic PPARγ plays opposite roles in rodent models and humans [37]. This assumption is supported by the lack of statistical correlation between PPARγ expression and the progression of NAFLD [38], and the anti-steatogenic effect of PPARγ exogenous agonists: thiazolidinediones (TZD) on NAFLD patients [39, 40]. It should be noted, however, that TZD activates PPARγ in multiple tissues to increase whole body insulin sensitivity, and their use is always associated with lifestyle modifications that could enhance the reduction of liver fat. By contrast, some research articles provide clear evidence that support an association between hepatic PPARγ expression (or the pathways controlled by PPARγ in the liver) and NAFLD/NASH in humans [10-12, 41-43]. In a recent transcriptomic analysis performed by RNAseq, Hansen et al. [44] did not detect significant differences in hepatic PPARγ expression in patients with NASH (BMI 33.9 +/- 6.2 kg/m2, patients recruited in Aarhus, Denmark [45]) when they were compared with normal weight individuals (BMI 23.1 +/-1.6 kg/m2) recruited in Copenhagen, Denmark [45]. However, in a group of 102 obese patients with severe obesity (BMI 43.18 +/- 0.55) recruited in Murcia, Spain, for bariatric surgery, the expression of PPARγ was significantly increased in patients with NAFLD/NASH when compared to obese patients without NAFLD. Due to the diversity of genetic pools in humans, and the influence of ethnicity, metabolism, and social/economic factors in the development of NAFLD, further research is required to understand if PPARγ expression could be associated with NAFLD in obese humans. With the use of rodent models of NAFLD where hepatocyte-specific PPARγ expression can be knocked out, multiple reports clearly show that the expression of PPARγ increases in the liver of mice with NAFLD and contributes to the development of fatty liver disease [13, 19, 21, 22, 46]. In fact, we recently showed that not only does hepatocyte-specific PPARγ contribute to the progression of NASH in mice fed a HFCF diet for 24 weeks, but it also reduces the potential therapeutic effect of PPARγ agonists in the treatment of NAFLD/NASH [13], and promotes steatosis in mice with severe diet-induced obesity [14]. However, since the simultaneous induction of obesity and human-like NASH through the use of diets is a challenge in mouse models [24, 47], it remains to be tested if the expression of PPARγ in hepatocytes in obese mice contributes to the progression of NAFLD.

The knockout of a hepatocyte-specific gene in adult mice represents a significant advantage to assess the relevance of that gene after a metabolic dysfunction has been established in an experimental model. This approach has the potential to unveil new roles of the studied gene which could be masked by developmental compensation induced by the knockout. Here, we used adeno-associated virus (AAV) to selectively, and efficiently [48], knockout PPARγ in hepatocytes in mice with pre-established diet-induced obesity. Also, we fed adult (>10 weeks of age) mice, instead of peripuberal mice (∼ 6 weeks), with a HFD to promote metabolic dysfunction and to prevent an increase of lean mass by HFD [49]. This study confirms again that the bulk expression of hepatic PPARγ in lean and obese mice is derived from hepatocytes, and that high fat diets increase the expression of PPARγ in the liver, specifically that of PPARγ2 [14, 20, 50]. Both PPARγ isoforms bind the PPAR response elements and may have the same gene targets, as noted by the steatogenic effects of an adenovirus-mediated cytomegalovirus promoter- driven PPARγ1 in the liver of mice [36], or overexpression of PPARγ2 in AML-12 cells or mice [51, 52]. In part, due to this strong steatogenic effect of PPARγ in the rodent livers, it is odd that without a direct confirmation, an opposite effect of hepatocyte-specific PPARγ in human and mice is accepted. Taken together, our current data show that PPARγ2 exclusively increases in hepatocytes of mice fed a HFD which consequently upregulates steatogenic mechanisms that promote steatosis [14, 16, 51, 52]. The results of this study are highly relevant because in a cohort of obese patients (BMI 43.18 +/- 0.55), NAFLD/NASH was identified in 72 patients (70% of the cohort) which were associated with an increased expression of PPARγ and insulin resistance. However, it is not clear if the presence of PPARγ in hepatocytes promotes NAFLD/NASH by regulating peripheral metabolism indirectly, or with the regulation of intrahepatic mechanisms.

Our study clearly shows that even 34 weeks of HFD (60% Kcal from fat) is not enough to increase inflammation and fibrosis in the liver. However, in diet-induced (18 weeks of HFD) obese mice, only 16 weeks of HFCF diet promoted NASH in both male and female mice. The progression of HFCF-induced NASH in female mice is not extensively explored. Nonetheless, it was recently reported that long-term feeding (>30 weeks) with NASH- inducing diets promotes NASH with fibrosis in female mice, but at a lesser level than males [53, 54]. We also show that NASH is more accentuated in males than females with pre-established diet-induced obesity, with a significant staining of collagen fibers in male mice just after 16 weeks of HFCF diet. In addition, Pparg^ΔHep^ mice reveal a sex- dependent effect of HFCF diet on the transcriptome and metabolome that is associated with reduced regulation of hepatic PPARγ-targets and the progression of NASH in female mice as compared to the effect in male mice. While women have less prevalence of NAFLD than men this risk significantly increases in women after menopause [55]. Thereby, estrogen signaling may ameliorate the development of NASH in females by affecting peripheral metabolic dysfunction, and/or by reducing the effects of PPARγ in female liver. Overall, our study shows that although the knockout of PPARγ is effective in female mice, this knockout has a limited impact on the expression of hepatic PPARγ-target genes and the liver transcriptome and metabolome, as compared to that of male mice. Therefore, there is a potential crosstalk between estrogen receptor and PPARγ signaling in hepatocytes that could determine the progression of NAFLD.

Although the Amylin liver NASH (AMLN) diet or the Gubra AMLN (GAN) diet required more than 30 weeks to induced NASH in male mice [26, 44], these diets induce NASH after 16 weeks of feeding in leptin-deficient hyperphagic mouse model (ob/ob) [26, 56]. It should be noted that long term feeding of AMLN and GAN diets does not promote adiposity as 16 weeks of HFD-feeding [24, 53], despite the GAN diet increasing fat mass tissue [44]. This evidence suggests that inducing obesity and human-like NASH together in mice is a challenge. Indeed, here we report that HFCF diet reduces adiposity in obese mice. This effect of HFCF diet could be due to an increase in energy expenditure, or the reduced energy content of the HFCF (4.5 kcal/g vs 5.2 kcal/g of HFD) diet [13, 24, 47]. However, when the fat source of this HFCF diet (partially hydrogenated corn oil) is replaced with lard or palm oil, the mice show an increased amount of fat mass [24, 44], so, it is possible that the trans-fat reduces adiposity in mice. Also, glucose intolerance and insulin resistance are commonly associated with NAFLD and obesity, but long term-HFCF feeding did not promote glucose intolerance nor insulin resistance [13, 24, 53]. When mice fed AMLN or GAN diets are compared to chow-fed mice, these diets show significant negative effects on glucose homeostasis [24, 44, 56]. However, caution needs to be taken when chow diet is the control diet, since sources of fiber, types of carbohydrates, and the composition of proteins in the control diet could improve glucose homeostasis. As such, it is critical to use nutrient-matched open formula diets as control diet [57]. Overall, our data suggest that HFCF diet does not enhance obesity nor metabolic dysfunction, and therefore the hepatic phenotype of HFCF-fed mice should be a consequence of direct actions of the diet on the liver rather than on peripheral metabolism. Our results also reveal the challenges of the NASH-inducing diets to reproduce the spectrum of metabolism abnormalities of patients with NAFLD/NASH, and the need to develop new diets that induce human-like NASH in short periods of time.

Loss of hepatocyte PPARγ did not reduce steatosis in HFCF-fed mice. However, Pparg^ΔHep^ reduced fibrosis in male mice, and the expression of fibrosis-related genes in male and female mice. Our results challenge the well- known steatogenic effect of PPARγ in hepatocytes. However, it should be noted that an HFCF diet provides an elevated amount of cholesterol with the potential to become oxysterols and activate liver X receptors in hepatocytes [58]. The aforementioned is known to increase steatosis [59], and may overcome the protective effect of Pparg^ΔHep^ in the development of steatosis. Nonetheless, our data indicates a clear effect of Pparg^ΔHep^ in gene expression and metabolite levels. In addition, our observations are aligned with the potential pro-fibrotic effect of hepatocyte PPARγ in different models of fatty liver disease [46, 60] including our previous report with HFCF-induced NASH and methionine and choline deficient mice [13, 15]. Since the development of fibrosis is not directly caused by hepatocytes, PPARγ might alter the intercommunication between hepatocytes and non-parenchymal cells to promote it. Our unbiased approaches indicated that hepatocyte PPARγ could be associated with the regulation of methionine metabolism, as we previously reported [13], and with the levels of Hcy which are significantly elevated in control male mice fed a HFCF diet. Of note, hepatocytes metabolize most of dietary methionine and are a main source of Hcy. Briefly, Hcy is produced after demethylation of methionine in hepatocytes and it can be converted to glutathione. Also, hepatocytes can re-methylate Hcy into methionine via BHMT to prevent its toxicity. Of note, BHMT is the main methyltransferase expressed in hepatocytes and uses betaine as a methyl donor to recycle Hcy into methionine [61]. If the levels of Hcy increase in hepatocytes, it can be exported out of the cell and activate non- parenchymal cells. In fact, Hcy levels in plasma are strongly associated with NAFLD in humans [62, 63], and they could activate hepatic stellate cells and promote fibrosis [64]. In HFCF-fed mice with NASH, the elevation in Hcy levels is the result of impaired re-methylation [65]. Thereby, treatments with methyl donors, such as folate, cobalamin, and betaine that enhance re-methylation of Hcy, have been considered in different studies to reduce some of the toxicity caused by excess Hcy and to reduce fatty liver disease [66, 67]. The fact that Pparg^ΔHep^ increased of BHMT expression in the liver of HFCF-fed mice and was associated with a reduction in Hcy levels highlight the potential relevance of PPARγ in the control of methionine metabolism and the prevention of Hcy-related hepatoxicity. The understanding of the mechanism that PPARγ uses to negatively regulate BHMT expression and/or activity to promote NAFLD requires further investigation.

Overall, our study highlights the relevance of hepatocyte-specific PPARγ in the progression of NAFLD and NASH by generating Pparg^ΔHep^ in adult male and female mice with pre-established diet-induced obesity. This is supported by evidence showing an increased expression of PPARγ in a cohort of patients with severe obesity and NAFLD/NASH. Finally, our study suggests that contribution of hepatocyte PPARγ in the context of NAFLD / NASH progression is influenced by gender where it could be associated with the development of fibrosis in males.

## Methods

### Subject study design

A total of 102 consecutive and prospective bariatric patients were enrolled at the Virgen de la Arrixaca University Hospital (Murcia, Spain). Inclusion criteria were age range of 18–65 years and obesity of more than 5 years of duration with a body mass index (BMI) ≥40 kg/m^2^ or ≥35 kg/m^2^ with significant obesity-related comorbidities. Exclusion criteria were set based on evidence of other causes of liver disease, including viral hepatitis, medication-related disorders, autoimmune disease, hepatocellular carcinoma, haemocromatosis, Wilson’s disease, familial/genetic causes, or a previous history of excessive alcohol use (>30 g daily for men and >20 g daily for women) or treatment with any drugs potentially causing steatosis, such as tamoxifen, amiodarone, and valproic acid. The study was performed in agreement with the Declaration of Helsinki according to local and national laws and approved by the Ethics and Clinical Research Committees of the Virgen de la Arrixaca University Hospital (#2020-2-4-HCUVA).

Blood samples were collected from study patients after an overnight fast of at least 12 h and serum was separated by centrifugation. The levels of HbA1c, glucose, insulin, and ALT were measured with glycohemoglobin analyzer HLC®-723G8 (Tosoh Bioscience), and the Cobas Analyzer c702 and e801 (Roche). HOMA-IR was calculated as insulin (µU/ml) x glucose (nmol/L)/22.5. Intraoperative wedge liver biopsies from bariatric surgery patients at least 1 cm in depth were obtained. One section of the liver biopsy was snap-frozen and stored at -80°C for RNA isolation and the other was formalin-fixed and paraffin-embedded for histological assessment.

### Mouse study design

All mouse studies were approved by the institutional Animal Care and Use Committee of the University of Illinois at Chicago. PPARγ floxed breeders with a C57B1/6J background were originally purchased from Jackson Laboratories (Strain 004584, B3.129-Ppargtm2Rev/J, Bar Harbor, ME), and bred as homozygotes in a temperature- (22-24C) and humidity-controlled specific-pathogen free barrier facility with 14h light /10h dark cycle (lights on at 0600h). Male and female PPARγ floxed mice were maintained on a standard chow diet and, when they were 10 weeks-old, were switched to a LFD containing 10% kcal from fat (Cat # D12450J, Research Diets, Inc, New Brunswick, NJ) or a HFD containing 60% kcal from fat (Cat # D12492, Research Diets, Inc). After 16 weeks of LFD/HFD, adult-onset hepatocyte-specific PPARγ knockout (*Pparg*^ΔHep^) and control mice were generated with AAV: AAV8-TBGp-Cre (#107787-AAV8) and AAV8-TBTp-Null (#105536-AAV8, Addgene, Watertown, MA), respectively, as reported previously [13]. Two weeks later, half of the HFD-fed mice in each group (control and *Pparg*^ΔHep^) were switched to a diet containing 40% kcal from fat (partially hydrogenated corn oil), 2% cholesterol (w/w), and 22% fructose (w/w) (HFCF diet, Cat #D16010101, Research Diets, Inc) for 16 additional weeks. HFCF diet induces NASH after 24 weeks in lean mice [13, 25]. This diet has a formula similar to the original AMLN diet [26] where trans-fat has been replaced with partially hydrogenated corn-oil. The composition of the diets is provided in the **Supplemental Table S1**. After 34 weeks of special diets, all mice were euthanized by decapitation after a 4 h food withdrawal at 0700h.

As previously published by our group [13], we assessed whole body composition (fat, lean, and fluid) with a minispec LF50 body composition analyzer (Bruker, Billerica, MA), performed GTT, ITT, and collected blood and tissues after 4h food withdrawal at 0700h. Blood was collected in EDTA-coated microtainers from lateral tail vein bleeding or from trunk blood at sacrifice. Tissues were weighed and fixed in formalin or snap-frozen in liquid nitrogen for molecular analyses.

### Liver lipid extraction and plasma endpoint

Hepatic TG were extracted using isopropanol as previously reported [27]. Plasma NEFA, plasma and hepatic cholesterol and TG (Fujifilm Wako Diagnostics, Richmond, VA), and plasma ALT (Point Scientfic, Canton, MI) were all measured using colorimetric assays. Plasma insulin levels were measured using an ELISA kit (Mercordia, Uppsala, Sweden).

### Histology

Paraffin-embedded 5µm sections of human liver biopsies were stained for H&E, Masson trichrome, Periodic acid–Schiff, Perls and reticulin staining. All biopsies were reviewed and scored by trained liver pathologists from the Virgen de la Arrixaca University Hospital to determine the steatosis, activity, and fibrosis classification system (SAF) [28]. SAF assesses steatosis (0-3), activity (0-4): hepatocellular ballooning (0-2) plus lobular inflammation (0-2), and fibrosis (0-4). Patients with steatosis score = 0 did not have NAFLD. Patients with steatosis score ≥1 had NAFLD, and of these NAFLD patients, those with scores ≥1 for hepatocellular ballooning and inflammation had NASH.

Paraffin-embedded 5µm sections of mouse livers were stained with H&E to determine the NAFLD activity score (NAS) using the Kleiner system [29] in a blind manner by two different groups. NAS assesses steatosis (0- 3), inflammation (0-3), and ballooning (0-2). A NAS >5 was considered NASH, while NAS ≤5 was not considered NASH. Also, liver sections were stained with picrosirius-red and fast green counterstain as previously described [13] to determine the area of fibrosis with ImageJ [30] using a specific macro (**Supplemental Table S2**), and to determine the type of fibrosis in a blind manner by a clinical pathologist. Fibrosis was scored based on the location of stained fibers of collagen into the hepatic lobule: no fibrosis (0), perisinusoidal or periportal fibrosis (1), perisinusoidal and periportal fibrosis (2), bridging fibrosis (3), and cirrhosis (4).

### Hepatic gene expression: quantitative PCR analysis and RNAseq

Total RNA from human liver biopsies (10-20 mg) was extracted using Trizol extraction (Life Technologies, Carlsbad, CA) followed by a purification step using the GeneJET RNA Purification kit (Thermo Scientific). 1000 ng of total RNA were reverse transcribed using the High-Capacity RNA-to-cDNA kit (Applied Biosystems). Real- time PCR amplification was performed using the Power SYBR Green Master Mix (Applied Biosystems). Relative expression levels were analyzed using the 2-ΔΔCT method and 18S rRNA was used as housekeeping gene. Total RNA from mouse livers was extracted using Trizol extraction and used for RNAseq or reversed transcribed for quantitative PCR as previously described [31, 32]. The sequence of primers used in this study can be found in **Supplemental Table S3**. cDNA library preparation, sequencing, and bioinformatics analysis of RNAseq was performed by Novogene (Novogene Inc, Sacramento CA) [13]. The raw files and the processed files with the differential expressed genes (DEG) are published in the Gene Expression Omnibus (GEO) with the accession # GSE200352.

### Hepatic protein expression: Western-blot

Proteins were extracted and analyzed as previously described [14]. Protein membranes were incubated with either PPARγ antibody (C26H12) #2435 (Cell Signaling Technologies, Danvers, MA) at 1:1000 or BHMT antibody #ab96415 (Abcam, Waltham, MA) at 1:2000 overnight at 4°C. The blots were developed with Clarity Max Western ECL substrate (Bio-Rad, Hercules, CA) and analyzed using a ChemiDoc MP imager (Bio-Rad).

### Steady-State Metabolomics

20 mg of frozen liver was homogenized in 80% ultrapure HPLC grade methanol (Fisher Scientific) in ultrapure water on dry ice at 20 ul/mg. After an overnight incubation at -80°C the samples were vortexed rigorously for 60 seconds and centrifuged at 20,000 xg for 30 minutes at 4°C. The supernatant was transferred to new tubes and dried with a nitrogen evaporator. Dried metabolites were resuspended in 50% liquid chromatography-mass spectrometry-grade acetonitrile at a final concentration of 20 μg protein/μl and processed by Metabolomics Core Facility at Robert H. Lurie Comprehensive Cancer Center of Northwestern University as previously described [13]. Analysis of the identified hydrophilic metabolites was performed with MetaboAnalyst 5.0 software [33].

### Statistical analysis

Data was analyzed with GraphPad Prism (GraphPad, San Diego, CA). To analyze the data of patients, we performed Student’s t-test to assess the differences between groups. Outliers were identified in the gene expression data using the ROUT method of identifying outliers. To analyze the mouse studies, two-way analysis of variance (ANOVA) with post-hoc Tukey multiple comparisons test was performed for all analyses. Group comparisons were performed between either LFD- and HFD-fed mice, or HFD- and HFCF-fed mice for both control and Pparg^ΔHep^ mice. Data shown in graphs were presented as mean ± standard error of the mean (SEM). P values <0.05 were considered significant. All authors had access to the study data and had reviewed and approved the final manuscript.

## Acknowledgements

We thank Angelie Bacon, Danielle Pins, Dr. Gregory Norris for their technical assistance. Fixed samples and H&E-stained slides were processed by the Research Histology Core at the University of Illinois at Chicago. Metabolomics services were performed by the Metabolomics Core Facility at Robert H. Lurie Comprehensive Cancer Center of Northwestern University.

## Notes

**Conflict of interest:** Authors do not have any conflict of interest.

### Competing Interest Statement

The authors have declared no competing interest.

